# Diagnosing Neurodegenerative Diseases by Label-Free 3-D Imaging of Intestinal Samples

**DOI:** 10.64898/2026.06.15.732258

**Authors:** Doriane Hazart, Marwa Moulzir, Brigitte Delhomme, Pascal Derkinderen, Malvyne Rolli-Derkinderen, François Cossais, Peter H. Neckel, Hervé Suaudeau, Fabrice Licata, Martin Oheim, Clément Ricard

**Affiliations:** Université Paris Cité, SPPIN, Saints-Pères Paris Institute for the Neurosciences, CNRS, 45 rue des Saints Pères, F-75006 Paris, France; Nantes Université, CHU Nantes, INSERM, The Enteric Nervous System in Gut and Brain Disorders, F-44000 Nantes, France; Université Rouen Normandie, INSERM, Normandie Univ, ADEN UMR 1073 Nutrition, Inflammation and Microbiota-Gut-Brain Axis, F-76000 Rouen, France; Institute of Clinical Anatomy and Cell Analysis, University of Tübingen, D-72074 Tübingen, Germany; Service de Microscopie Commun (SCM), BioMedTech Facilities, CNRS UAR 2009, INSERM US 36, Université Paris Cité, F-75006 Paris, France

**Keywords:** Parkinson’s disease, Enteric Nervous System, Neuropathology, Autofluorescence signals, Label-free imaging, Confocal microscopy, Diagnosis

## Abstract

Early diagnosis of Parkinson’s disease (PD) remains challenging because motor symptoms appear only after extensive neurodegeneration, and a definitive diagnosis still relies on *post-mortem* neuropathology. Increasing evidence implicates the enteric nervous system (ENS) in prodromal disease stages, but routine ENS-based diagnosis is limited by the complexity of intestinal tissue organization and the need for specific labeling strategies. Here, we developed a label-free autofluorescence (AF) imaging workflow combined with unbiased morphometric analysis to identify neurodegenerative alterations in fixed human colonic tissue. Using a correlative multiscale imaging approach, we generated a database of almost 800 high-resolution confocal images from myenteric and submucosal plexuses of controls, PD, and Alzheimer’s disease (AD) patients. Blind evaluation by four expert histologists showed reliable identification of control tissue but lower sensitivity for pathological cases, reflecting the heterogeneous distribution of ENS lesions. Semi-quantitative and morphometric image analyses identified a distinct population of enlarged enteric neurons, termed large neural cells (LNCs), strongly enriched in PD and AD compared with controls. LNCs contained autofluorescent cytoplasmic inclusions and frequently prominent nucleoli, both features largely absent from control tissue independent of aging. Co-localization with the amyloid-binding probe Amytracker (AmyT) demonstrated that AF granules correspond to β-sheet-rich protein aggregates rather than merely age-related lipofuscin granules. Similar alterations were detected in intact three-dimensional (3-D) colonic biopsies, demonstrating the feasibility of volumetric ENS imaging without tissue clearing. Together, our results establish label-free AF imaging as a rapid and clinically compatible strategy for detecting enteric neurodegenerative pathology. This approach provides a framework for the future development of ENS-based biomarkers and supports the use of volumetric intestinal imaging for early diagnosis of neurodegenerative diseases.

## Introduction

Parkinson’s disease (PD) is a progressive neurodegenerative disorder characterized by bradykinesia, rigidity, and resting tremor. Diagnosis currently relies on clinical evaluation of these motor symptoms according to criteria established by the Movement Disorder Society [3, 4, 39]. However, by the time of diagnosis, about 50–70% of dopaminergic neurons in the substantia nigra pars compacta have already degenerated [15, 22], indicating that the disease is already advanced.

Distinguishing idiopathic PD from atypical parkinsonian syndromes also remains difficult, with post-mortem studies reporting diagnostic error rates of up to 25% [31]. At present, definitive confirmation of PD still relies on post-mortem detection of Lewy bodies (LBs) in the substantia nigra pars compacta [16], which remains the histopathological gold standard.

Beyond motor symptoms, PD is associated with non-motor manifestations such as anosmia, REM sleep disorders, constipation, anxiety, and depression, which may appear years or even decades before motor onset [39]. This prodromal phase has driven major efforts to identify early biomarkers for diagnosis and the development of neuroprotective therapies.

Among emerging approaches, the detection of pathological α-synuclein aggregates has become a major focus of biomarker research in PD and related synucleinopathies [12, 19, 24, 35]. Techniques such as RT-QuIC (real-time quaking-induced conversion), adapted from prion disease research, can detect α-synuclein aggregates in biological samples including cerebrospinal fluid, saliva, and skin biopsies [17, 20, 42, 51]. Other strategies under investigation include inflammatory proteins, microRNAs, blood metabolites, gut microbiota analysis, and functional or metabolic MRI [5].

Constipation is a well-known early hallmark of PD that can precede motor symptoms by years [1, 11, 21, 33, 48]. This has increased interest in the enteric nervous system (ENS), a large network of about 600 million neurons that autonomously regulates gastrointestinal functions while communicating with peripheral and central pathways. The ENS is organized into two main layers: the myenteric (Auerbach’s) plexus, which controls gut motility, and the submucosal (Meissner’s) plexus, mainly involved in secretory and vascular functions [14, 23, 53].

Constipation and gut inflammation may not simply be consequences of PD. According to Braak’s hypothesis, disease could originate in the periphery, with pathological α-synuclein aggregates first appearing in the ENS, or the olfactory bulb, before spreading to the central nervous system via the vagus nerve or olfactory pathways, respectively [6–10, 30]. Supporting this idea, pathological α-synuclein aggregates have been detected in colonic biopsies from patients at early disease stages, making the ENS a potential source of prodromal biomarkers [13, 45]. However, early PD diagnosis based on ENS imaging remains challenging due to its complex three-dimensional (3-D) organization, the lack of standardized diagnostic criteria, and the difficulty of identifying neuronal ganglia in colonic biopsies.

In this work, we present a label-free 3-D imaging workflow combined with unbiased image analysis for PD diagnosis. Its relatively easy, speed and the absence of lengthy staining or antibody-labelling make our workflow potentially compatible with clinical practice. Using autofluorescence (AF) imaging of the ENS in fixed human colon tissue, first we identify neuronal and subcellular features associated with PD and other neurodegenerative diseases without the need for pathological labeling, enabling faster analysis.

It should be noted that the analyzed specimens corresponded to Lewy body disorders (LBD) and non-Lewy body disorders (non-LBD) cases rather than clinically defined PD and Alzheimer’s disease (AD) cases [26, 27]. For the sake of simplicity and readability, these groups are referred to throughout the manuscript as “PD” and “AD”, respectively.

We apply the method to controls, PD, and AD samples, perform a blind evaluation by expert histologists, and identify morphological features linked to accurate diagnosis. Finally, we extend the approach to intact 3-D colonic biopsies, highlighting the importance of volume imaging for future diagnosis of neurodegenerative diseases.

## Materials and Methods

### Ethics statement for human samples

Specimens of human descending colon have been obtained from the Neurological Tissue Bank of the Biobank-Hospital Clinic-IDIBAPS and described previously [26]; they were kindly provided by Dr. Ellen Gelpi. Clinicopathological diagnoses comprised LBD in 3 cases (including PD and dementia with LBs) and 3 cases off non-LBD (including 2 AD and one progressive supranuclear palsy, PSP). The control group comprised 10 individuals without neurodegenerative disorders who had colonic resection for colorectal cancer, diverticulitis and Crohn’s disease sampling was performed in macroscopically normal segments of uninvolved resection margins). Additional demographic and neuropathological features are summarized in ***Fig.1a***.

For the IDIBAPS biobanking, the study was approved by the Institutional Ethics Committee of the Hospital Clinic de Barcelona (registry number 2007/3906 and 2013/8201) and written informed consent was obtained from donors and/or next of kind for obtaining postmortem brain and body tissue samples for diagnostic and research purposes. For additional controls, sampling and procedures were performed according to the guidelines of the French Ethics Committee for Research on Humans and registered under the no. DC-2008-402.

Additional colonic biopsies from 4 patients with PD were used for final method validation (ColoBioParker NCT01353183). The study protocol by the National Committee on Ethics and Human Research (Comité de Protection des Personnes Nord Ouest IV) and registered on ClinicalTrials.gov (identifier NCT04652843).

### Sample preparation

#### Deparaffinization of gut histological sections

Slides were first dipped 20 times in Histoclear™ (Sigma-Aldrich), incubated for 10 min in a second bath of Histoclear™ before being subsequently dipped 10 times in a third bath of the same solution.

Sections were progressively re-hydrated through successive passages in baths of decreasing concentrations of ethanol (95%, 70%, and 50%). They were then incubated during 10 min in tap water to complete rehydration.

After drying, sections were mounted in a medium containing 90% glycerol, 10% Tris 0.2 M and 25 g/l DABCO (1,4-Diazabicyclo[2,2,2]octane, triethylenediamine).

#### Amytracker staining

Slides previously observed by AF were disassembled, and the glycerol-based mounting medium was removed by rinsing in PBS 1X baths.

Slides were then incubated for 30 minutes at room temperature, protected from light, in a solution of Amytracker (AmyT) 630 (Ebba Biotech) diluted 1:1000 in PBS 1X.

Finally, slides were air-dried and remounted in glycerol.

### Tissue imaging

#### Slide scanner

Samples were imaged in AF integrally on a slide scanner (AxioScan Z1, Carl Zeiss, Oberkochen, Germany). Images were acquired upon 430nm excitation, and fluorescence was extracted with a TBS 450+538+610 dichroic (Zeiss) and collected between 450 nm and 750 nm with a TBP 467/24+555/25+687/145 emission filters (Zeiss) using a 20X /0.8NA Plan-Apochromat objective (Zeiss). The acquired wide-field AF images allowed us to first identify and tag the coordinates of all nerve ganglia of the myenteric and submucosal plexuses, respectively. Their spatial coordinates (x, y) were recorded on the slide scanner images and a custom FIJI macro used to map the coordinates onto the confocal images so as to re-image the very same Region of Interests (ROIs) on the LSM880 at increased resolution and detail.

#### Confocal microscopy

Higher-resolution images of previously identified ROIs of tissue sections and biopsies were acquired on an inverted image-scanning confocal microscope (LSM880 AiryScan, Carl Zeiss, Oberkochen, Germany) fitted with a 40X/1.2 NA Plan-Apochromat water immersion lens (Zeiss). AF images were excited at 405 nm and fluorescence collected through a MBS – 405 dichroic on the spectral detector between 417 nm and 740 nm. The confocal pinhole diameter was systematically set to 1 Airy unit.

#### Spectral confocal acquisition

In some cases, we further split the detected AF into 32 spectral channels. Spectral images were acquired upon 405-nm excitation at the level of myenteric’s and submucosal’s plexuses. For each plexus, at least 3 clearly identified AF granules were delineated, the spectra extracted and normalized and data from neurodegenerative diseases and control intestinal slices compared.

### Image analysis, quantification, and statistics

#### Blind Diagnostic Survey

A dataset of 787 AF images and an evaluation questionnaire were submitted to four expert histologists through a custom online interface specifically developed for this study. Each participant accessed the survey via a unique password, and images were presented in random order without clinical information other than the presence of control and PD samples. No prior training was provided, and participants could not revisit previously given answers, reproducing conditions close to routine diagnosis.

For each image, participants were asked to (i) identify the observed structure (myenteric’s plexus, submucosal’s plexus, or other), (ii) provide an evalution (PD, healthy, or other pathology), (iii) rate their confidence level, and (iv) optionally describe the morphological criteria supporting their decision (***Fig.SI1***).

Responses were automatically exported to Excel and merged into a single dataset for subsequent analysis of diagnostic performance and inter-observer robustness.

#### Semi-quantitative analysis

The same image dataset was independently analyzed by the first author using predefined morphological and quantitative descriptors, with knowledge of the sample identity. A separate online questionnaire was used to standardize evaluation. Criteria of AF analysis included (i) plexus type (myenteric plexus, submucosal plexus, or undetermined), (ii) presence of small neural cells (SNCs) or large neural cells (LNCs), (iii) presence of cytoplasmic granules and/or nucleoli, and (iv) degree of infiltrating cells. Semi-quantitative scales (none, a few, many) were applied to selected features, and a free-text field allowed to note additional observations ***(Fig.SI2)***.

#### Correlation of diagnostic responses with semi-quantitative descriptors

Expert histologists’ evaluation were matched image by image with the semi-quantitative descriptors to identify morphological features associated with neurodegenerative disease recognition. This analysis enabled assessment of diagnostic accuracy and correlation of responses with specific image features.

Although images from AD patients were included, AD was not proposed as a diagnostic option. Therefore, an AD image classified as “Parkinson’s” was considered correctly identified as pathological rather than healthy tissue.

Each image received a score corresponding to the number of expert histologists providing a correct diagnosis, ranging from 1 to 4.

#### Morphometric characterization of ENS ganglia cells

Putatively enlarged neurons were manually identified, delineated and their area (µm²) and perimeter (µm) measured. In total, 2,016 cells were analyzed in this manner. We then plotted formal all cells their perimeter vs. Area, which allowed us to readily indentify LNCs – predominantly present in neuropathologic conditions and SNCs present in all samples. Morphometric measurements were performed using ImageJ/FIJI [44].

#### Co-localization of AF and Amytracker staining

Fluorescence was excited at 405 nm for AF and at 633 nm for AmyT and was collected between 415 nm and 600 nm for AF and between 640 nm and 758 nm for AmyT, respectively. The confocal pinhole diameter was again 1 Airy unit.

We analyzed overlap between AF and AmyT staining by plotting line profiles across AF granules labeled with AmyT. The measured intensities were normalized to their respective maximum.

#### Data processing

Image-derived measurements from the semi-quantitative analysis were processed using Microsoft Excel, and statistical analyses were performed using Prism (GraphPad, v10). Given the small sample size, all measurements are presented as full datasets, with the mean ± standard deviation (SD) overlaid. All experiments were conducted on n ≥ 3 independent individuals. Statistical comparisons were performed using the non-parametric Mann–Whitney test.

For the morphometric analysis, measurements of cell area (µm²) and perimeter (µm), obtained from a total of 2016 cells, were analyzed using Prism (GraphPad, v10). Data are presented as mean ± SD. Statistical comparisons between groups were performed using an unpaired t-test.

## Results

### Development of a large-scale autofluorescence imaging workflow

We hypothesized that high-resolution AF imaging and morphological analysis of ENS tissue could reveal cellular and subcellular alterations associated with neurodegenerative pathology.

Our study was carried out on a cohort of 16 subjects, divided into four distinct groups: young controls (age: 31.5 ± 11.8 years ; n = 4), aged controls (70.3 ± 10.7 years ; n = 6), patients with PD with a post-mortem diagnosis of LBD and the presence of α-synuclein deposits in the ENS [26], (81 years ± 4.6 ; n = 3), and patients diagnosed with AD and progressive supranuclear palsy without α-synuclein aggregates in the ENS (85.7 ± 7.8 years ; n = 3).

Control samples were obtained from patients without known neurodegenerative diseases and originated from healthy resection margins of surgically-removed colonic pieces from other pathological conditions ***(*Fig.1*)***.

**Fig. 1.**
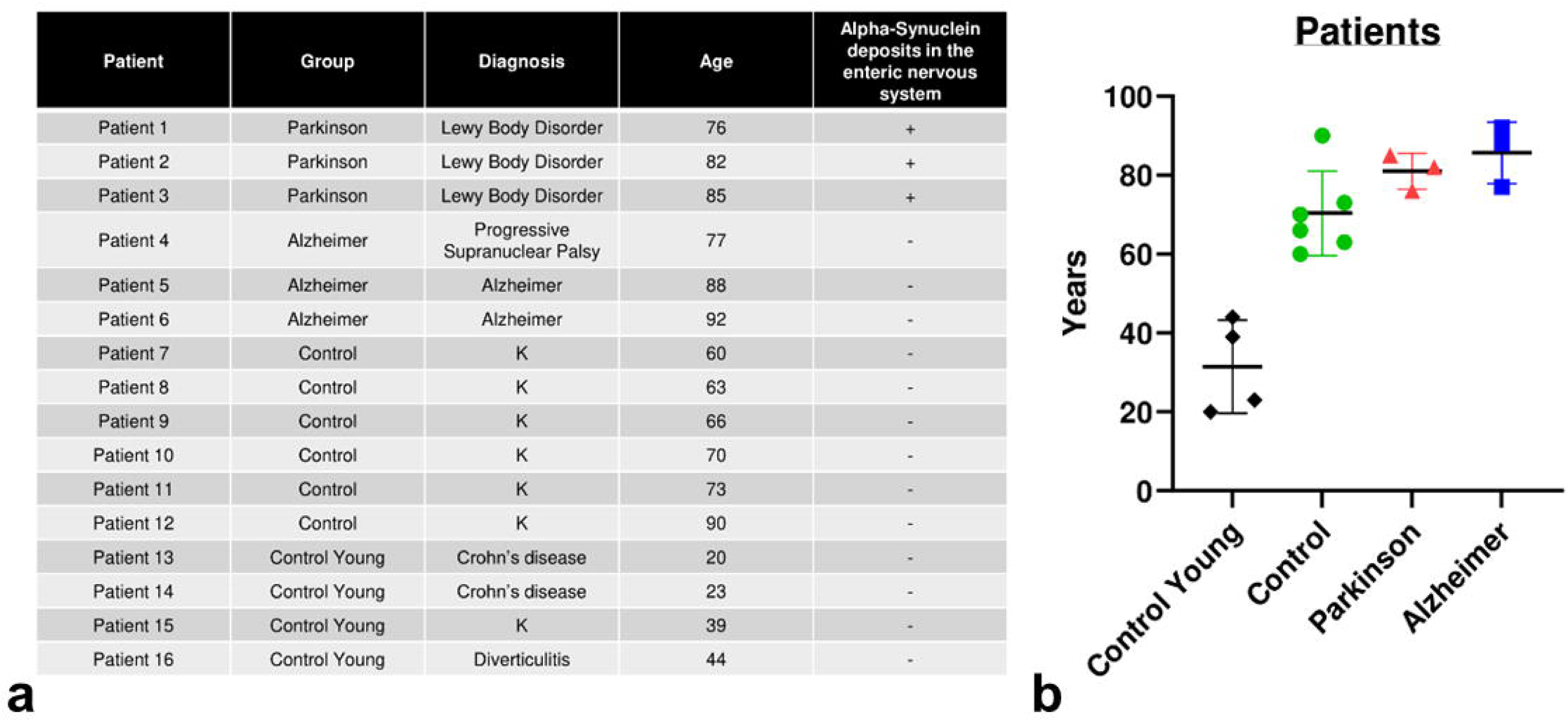
Definition of the cohort for the comparative study. a. Summary table of patients included in the study, categorized by group (Parkinson’s, Alzheimer’s, older and younger controls), diagnosis, age, and the presence or absence of α-synunclein deposits in the ENS; b. Age of patients by group: young controls (black), aged controls (green), Parkinson’s (red), Alzheimer’s (blue). Each point represents an individual. The bars correspond to the mean ± standard deviation

On each slide, we first located all enteric nerve ganglia. However, acquiring high-resolution scans of the entire tissue surface (about 6 cm^2^) using a confocal laser scanning microscope (CLSM) is impractical, because of the long acquisition times on a scanning-type microscope and the simple amount of image data generated. We therefore used a correlative, multi-scale AF imaging workflow instead, to first identify ROIs on overview images before zooming in at higher resolution on identified ROIs ***(*Fig.2*)***.

**Fig. 2.**
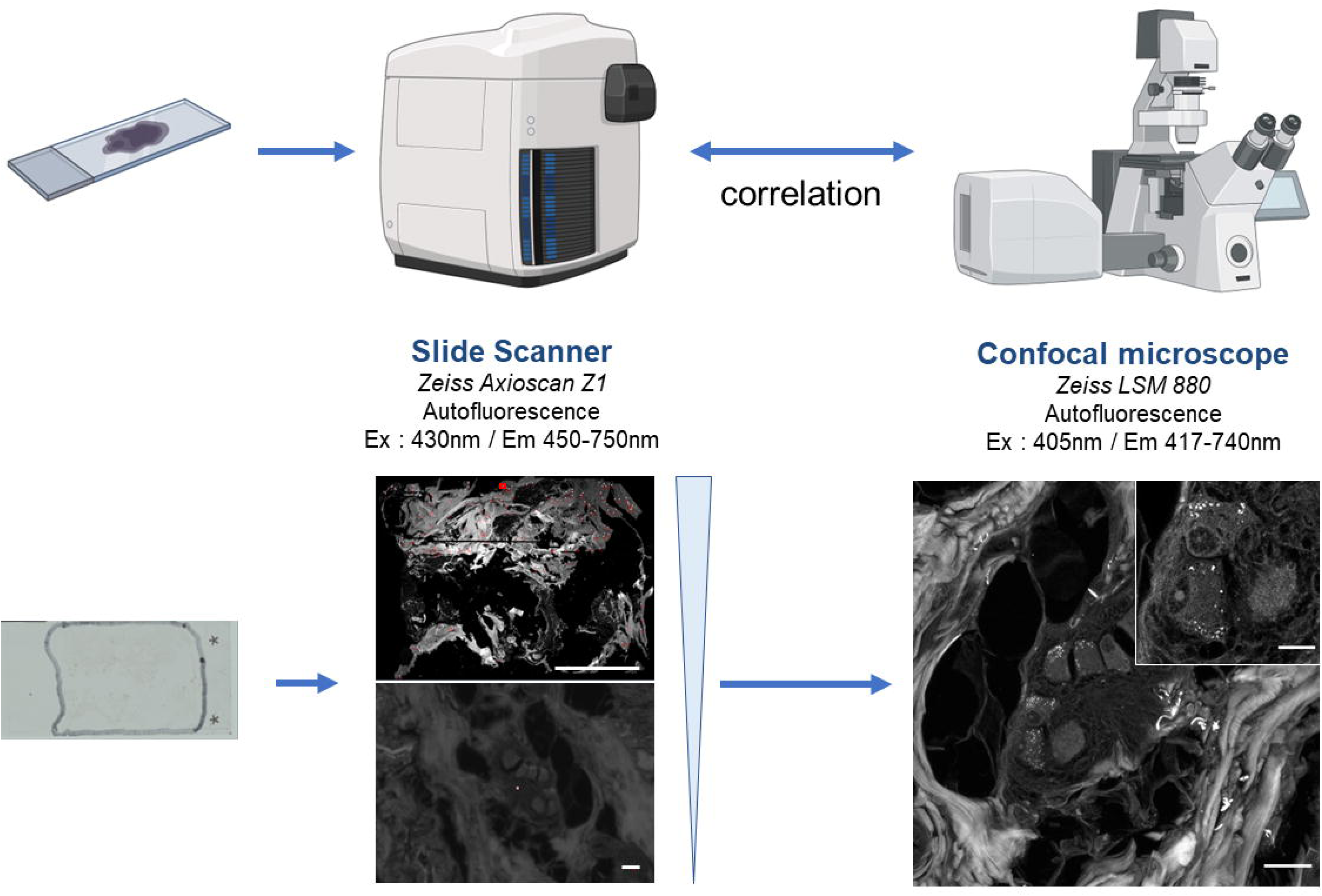
Correlative AF imaging workflow for morphological analysis of human ENS. Slide Scanner scale bars: 1cm (top), 25 µm (bottom); Confocal microscope scale bars: 25 µm *(created with BioRender)*

After de-paraffinization and mounting, a low-resolution (20X/0.8NA air) overview of the entire intestinal tissue slice was acquired on a robotic slide scanner ***(Fig.2, bottom left image)***. We then identified and located all ENS nerve ganglia on the slide. The specific (x, y) coordinates of each ROI were transferred to the stage command of the CSLM by aid of a custom macro. This coordinate transform allowed us to re-image only a subset of the slice at higher resolution (40X/1.2NA water) ***(Fig.2, bottom right image)***.

Applying this protocol yielded 787 diffraction-limited images (213 x 213 µm; pixel size = 208 nm) constituting an image database for qualitative analysis and detailed characterization of the human ENS both in control and neuropathological conditions.

### Towards early PD diagnosis: blind expert evaluation

To assess whether AF imaging of ENS ganglia can support PD diagnosis, four expert histologists independently and blindly analyzed the full image dataset. No clinical information was provided except that the cohort included control and PD cases. Each expert classified images as healthy, PD, or other pathology via an online questionnaire ***(Fig.SI1)***, with an optional free-text justification of the criteria used.

Overall performance varied across observers and conditions. For control samples, two expert histologists (A and B) achieved high accuracy (∼95–100%), while C and D showed lower agreement (∼55–60%). In contrast, PD and AD images were frequently misclassified as healthy by all observers. For PD cases, A and B correctly identified only ∼10% as PD, while C reached ∼30% accuracy and D ∼20%, with many cases distributed between “healthy” and “other pathology.” ***(Fig.3a)***.

**Fig. 3.**
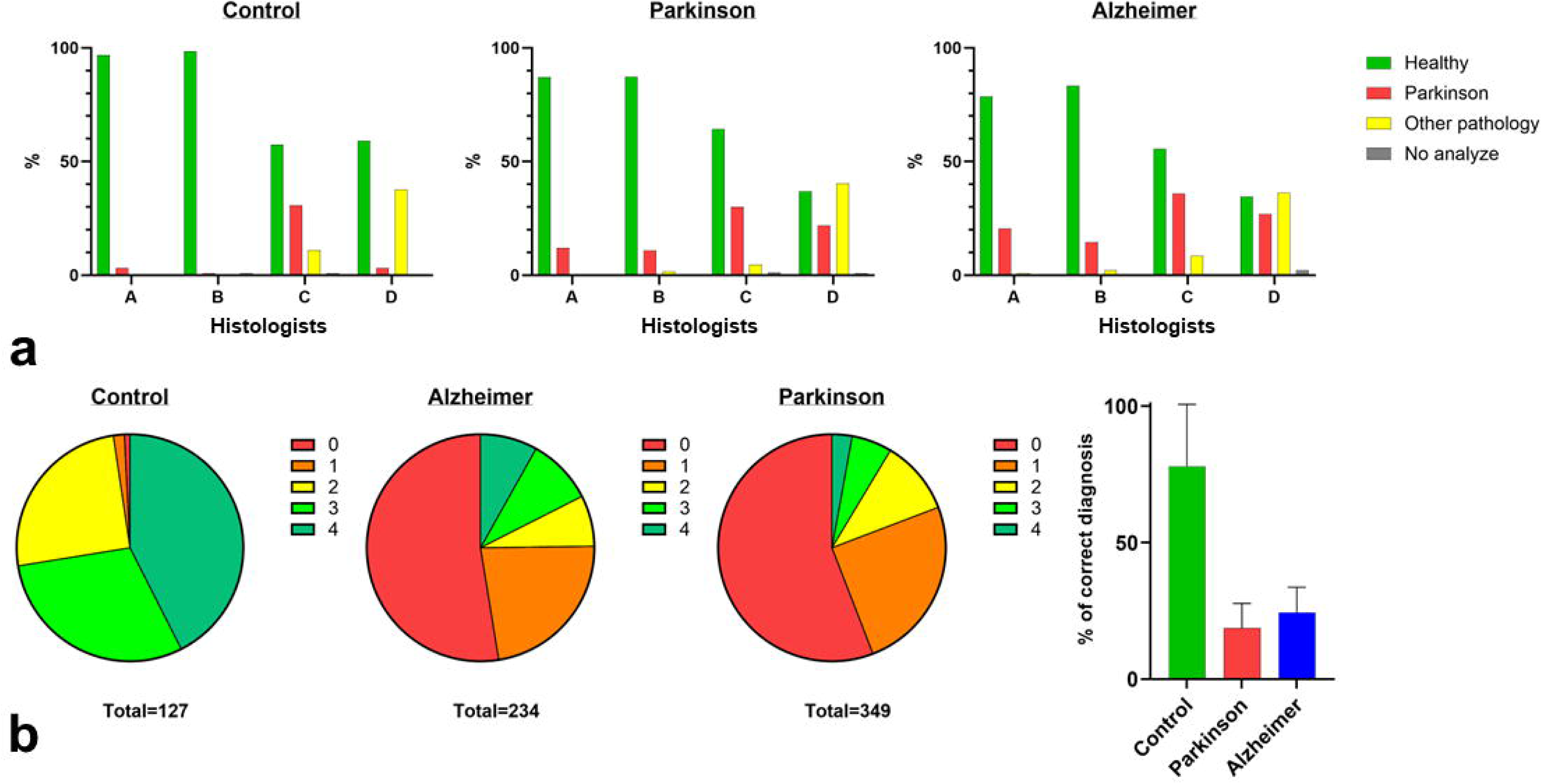
Diagnostic performance of expert histologists using fast autofluorescence imaging of ENS ganglia. a. Distribution of diagnostic classifications assigned independently by four expert histologists (A–D) across image categories (Control, Parkinson’s disease (PD), and Alzheimer’s disease (AD)). Results are expressed as percentages of images classified as healthy (green), PD (red), other pathology (yellow), or not analyzed (gray); b. Diagnostic performance assessed using a consensus score (0–4), corresponding to the number of expert histologists assigning the correct diagnosis (0: none; 4: full agreement). Pie charts illustrate the distribution of scores for each image category (total number of images indicated). The bar graph summarizes the overall percentage of correct diagnoses (mean ± SD)

### Quantitative assessment of diagnostic performance

Diagnostic responses were pooled across all expert histologists using a semi-quantitative score ranging from 0 (no correct classification) to 4 (full consensus correct diagnosis). Control images were generally well recognized, with >75% reaching high agreement scores (3–4) and being correctly identified as healthy. In contrast, pathological samples (PD and AD) showed low concordance, with <25% achieving high agreement scores and most falling in the 0–2 range, reflecting misclassification or lack of consensus ***(Fig.3b)***. Overall, control images were identified with moderate to high confidence, whereas pathological cases showed poor diagnostic consistency. This suggests that ENS-associated neurodegenerative features may be heterogeneous or variably detectable across thin ENS tissue slices.

As AD was not a diagnostic option and was unknown to the observers, correctly identified AD images were counted as “PD,” reflecting recognition of neuropathology rather than disease-specific classification.

Collectively, our findings highlight both the potential of AF imaging to detect the absence of ENS abnormalities and the need for more robust criteria to reliably distinguish healthy tissue and differentiate between neurodegenerative conditions.

### Qualitative, morphometric and semi-quantitative analysis of ENS sections

To better understand the criteria that motivated the experts’ evaluations, we analyzed the free-text responses from the blind survey. However, these comments provided little consistent information, with the only recurrent feature mentioned being “lipofuscin granules”.

We therefore performed a complementary qualitative, morphometric, and semi-quantitative analysis. From 787 images, 741 corresponding to ganglia of the myenteric (355) and submucosal (386) plexuses were manually examined and search for stereotypical features associated with control, PD, and AD samples.

Control tissues showed a regular, homogeneous neuronal organization with rounded cell bodies and no evident cytoplasmic abnormalities ***(Fig.4a)***. In contrast, PD and AD samples displayed marked structural heterogeneity, including the presence of markedly enlarged neurons, which we termed Large Neural Cells (LNCs). These LNCs could further be grouped into three subtypes: (i) LNCs without detectable inclusions, (ii) LNCs containing autofluorescent cytoplasmic granules, and (iii) LNCs showing both granules and prominent nucleoli ***(Fig.4b)***. LNCs, particularly those containing cytoplasmic granules and/or nucleoli, were absent or extremely rare in control tissues, suggesting a potential link to neuropathology.

**Fig. 4.**
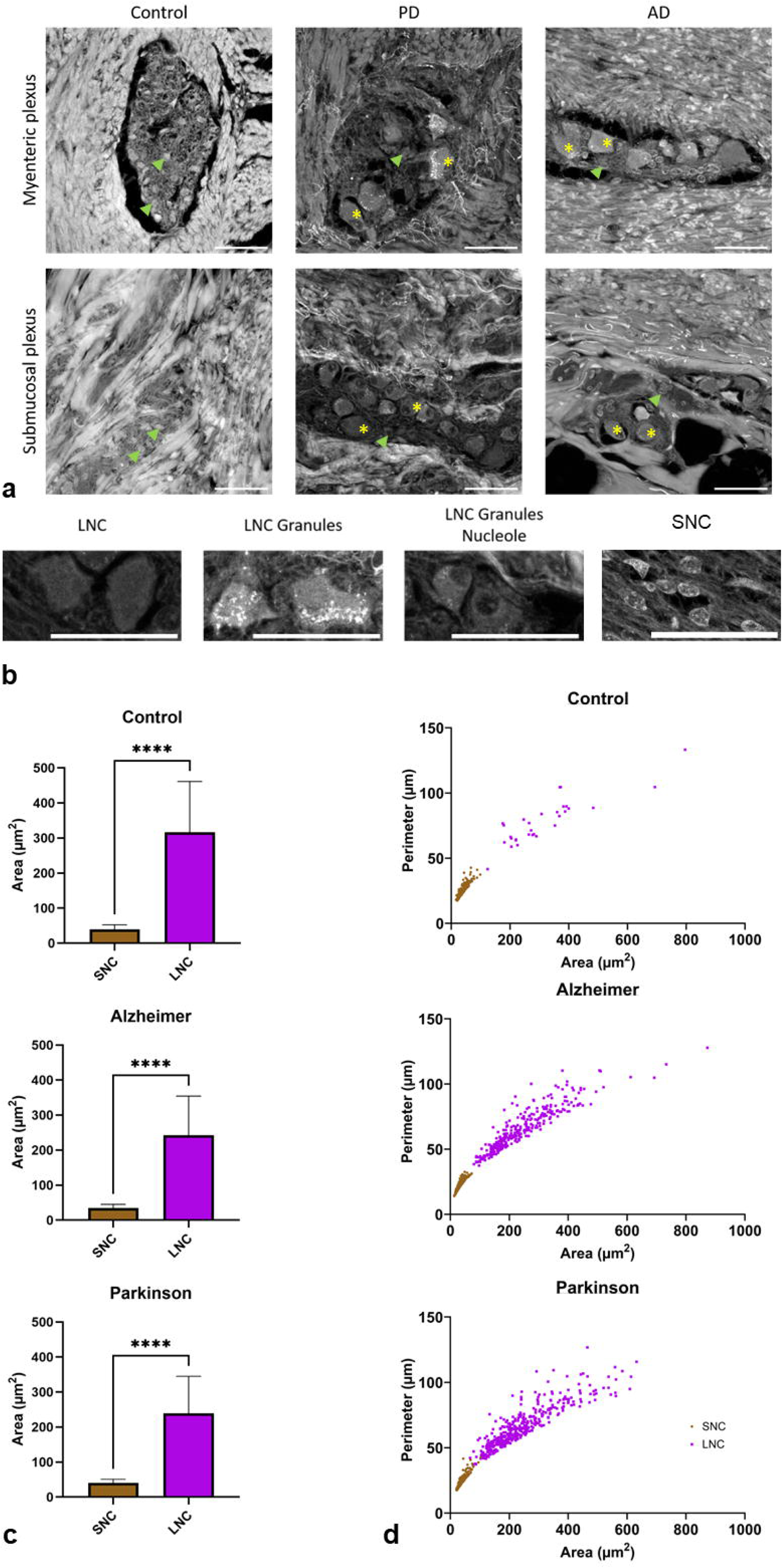
Qualitative and morphometric analysis of ENS cells. a. Representative AF images of nerve ganglia from the myenteric plexus (top row) and the submucosal plexus (bottom row) for the three groups studied: controls, patients with Parkinson’s disease (PD), and patients with Alzheimer’s disease (AD). Green arrows show SNCs and yellow asterisks show LNCs; b. Representative images illustrating the different cell types (LNCs and SNCs), Scale bars: 50 µm; c. Bar graphs show comparison of mean area (µm²) between SNC (brown) and LNC (purple) populations across the three conditions (statistical test is a t-test with **** p < 0.0001); d. Scatter plots show the relationship between perimeter (µm) and area (µm²) for individual SNC (brown) and LNC (purple) structures

In parallel, a second population of small neural cells (SNCs) was systematically observed ***(Fig.4b)*** in control and pathological samples, suggesting they likely represent a normal ENS cell population unaffected by disease.

Although the distinction between LNCs and SNCs was visually striking, we further substantiated our observation by quantitative morphometry, measuring the area and perimeter 760 LNCs and 1256 SNCs.

Across all groups (Control, AD, PD), LNCs were significantly larger than SNCs (244,1 ± 110,8 µm^2^ vs 38,02 ± 11,78 µm^2^) ***(Fig.4c)***.

Scatter plots of cell area versus perimeter further separated the two populations, with LNCs showing larger and more irregular morphologies characterized by disproportionately high perimeters ***(Fig.4d)***, lending further support to the existence of two distinct cell populations. We note that although LNCs were enriched in both PD and AD samples compared to controls, they did not allow discriminating between pathologies.

Building on these morphometric criteria, we quantified LNC populations in different conditions ***(Fig.5a)***. The proportion of images containing at least one LNC increased from 17% in controls to 40% in PD and AD (2.3-fold increase) ***(Fig.5a–c)***. This effect was even stronger for LNC subtypes with granules (38% vs 10.5%, 3.2-fold) and with granules plus nucleolus (20% vs 2%, 10-fold). Similar trends were observed in both the myenteric and submucosal plexuses ***(Fig.5c)***.

**Fig. 5.**
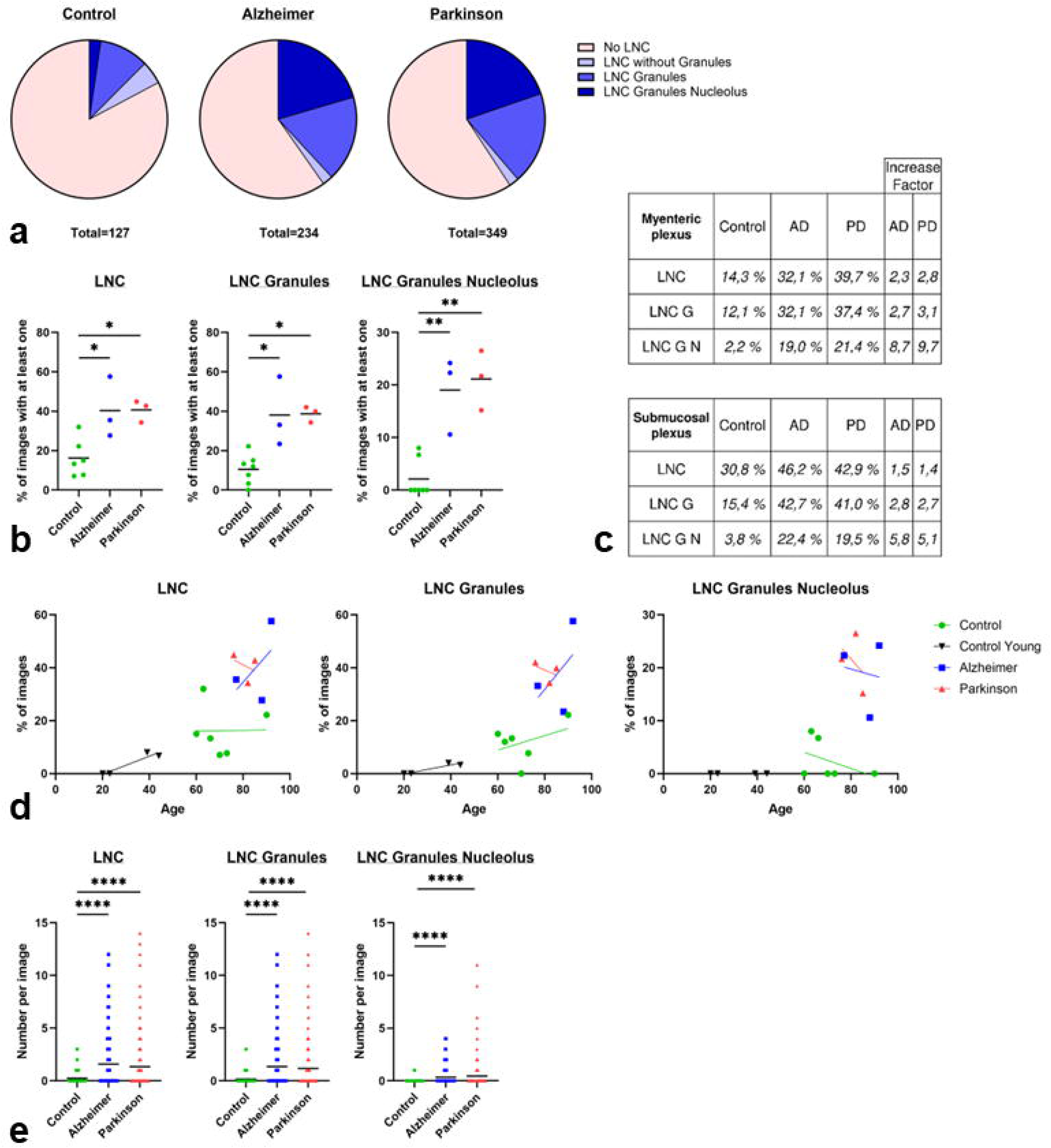
Statistical analysis of LNCs in the ENS of control, Alzheimer’s disease and Parkinson’s disease groups. a. Distribution of LNC types observed in samples from control, Parkinson’s, and Alzheimer’s groups. The sections represent, respectively, images without LNCs (pink), with LNCs without granules (purple), with LNCs and granules (light blue), and with LNCs, granules and nucleolus (dark blue). The total number of images analyzed is indicated below each graph; b. Percentage of images containing at least one LNC, LNC with granules, or LNC with granules and nucleolus in the different groups (control, Alzheimer’s, Parkinson’s). Each point represents an individual. The horizontal bars indicate the mean; c. Summary tables of LNC frequencies observed in the myenteric and submucosal plexuses, expressed as a percentage of positive images. The increase factors in the Alzheimer’s (AD) and Parkinson’s (PD) groups are indicated in the last column; d. Correlation between age and the percentage of images containing LNCs, LNCs with granules, and LNCs with granules and nucleoli. Groups are distinguished by color: black (control young), green (control), red (Parkinson), blue (Alzheimer); e. Mean number of different types of LNCs per image for each group. Each point represents one image. The bars indicate the mean. Each statistical test is a Mann-Whitney test with *p < 0.05, **p < 0.01, and ****p < 0.0001

Age alone could not account for these differences, as aged controls showed only limited increases in the number of images containing LNCs compared to young controls, far below those observed in pathological groups ***(Fig.5d)***. Moreover, the number of LNCs per image was consistently higher in disease samples across all subtypes ***(Fig.5e)***, supporting LNC accumulation as a robust marker of neurodegeneration.

Together, these results identify LNCs - particularly granule- and nucleolus-containing forms - as a morphological signature of degenerative neuropathology, and not simply attributable to age. In contrast, SNCs showed no significant differences between groups, appearing on most images across controls (67%), AD (59%), and PD (45%) ***(Fig.6a–d)***. SNC subtypes with granules or nucleoli were rare (<10%) and showed no group-dependent variation or age correlation ***(Fig.6e)***, suggesting that SNCs represent a stable, constitutive ENS population of cells unaffected by pathology.

**Fig. 6.**
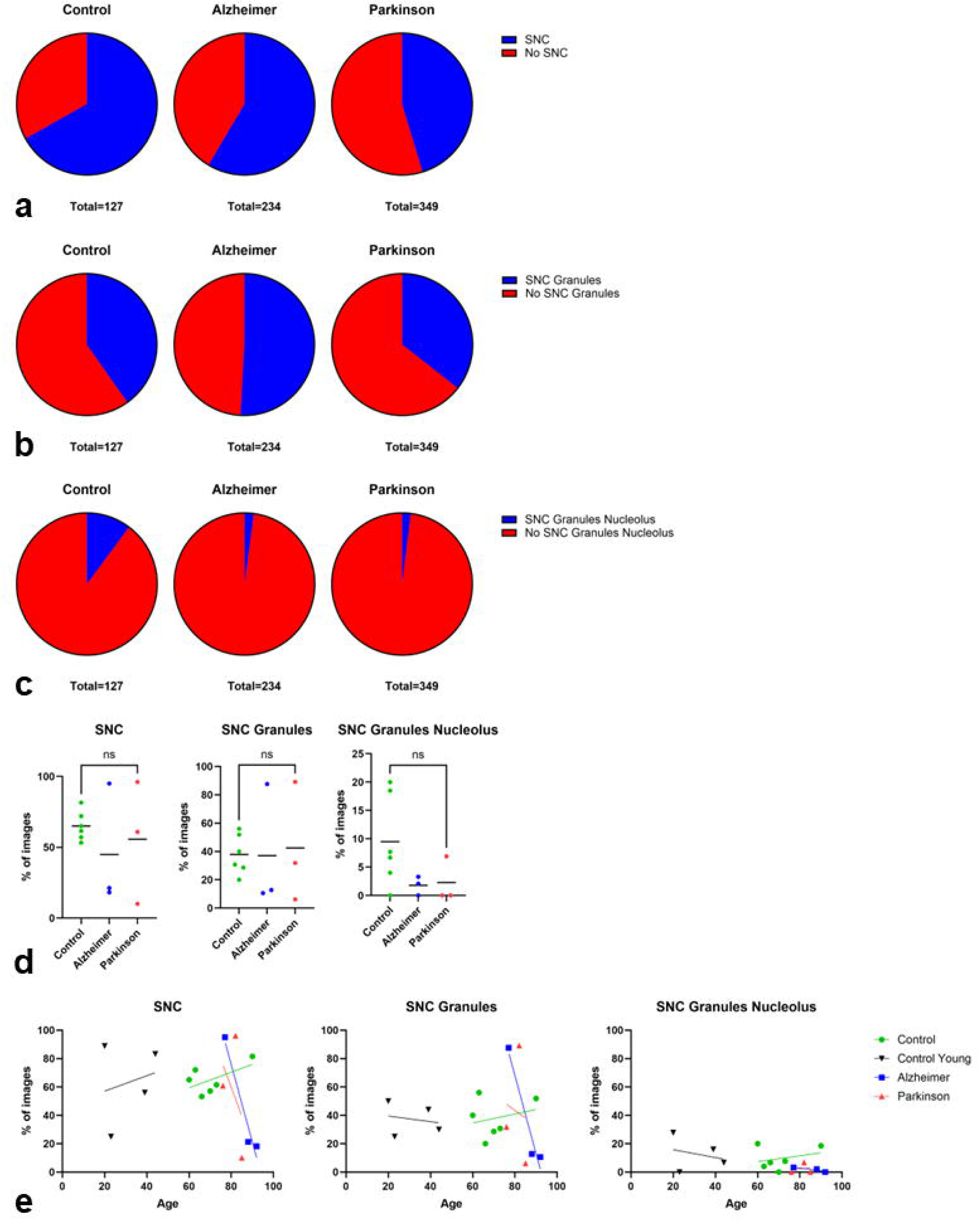
Statistical analysis of SNCs within the ENS in control patients, patients with Alzheimer’s disease, and patients with Parkinson’s disease. a. Distribution of images containing at least one SNC in the Control, Alzheimer’s, and Parkinson’s groups. The total number of images analyzed is indicated below each graph; b. Distribution of images containing at least one SNC with cytoplasmic granules. The total number of images analyzed is indicated below each graph; c. Distribution of images containing at least one SNC with cytoplasmic granules and nucleolus. The total number of images analyzed is indicated below each graph; d. Percentage of images containing at least one SNC, one SNC with granules, or one SNC with granules and nucleolus in the different groups (control, Alzheimer’s, Parkinson’s). Each point represents an individual. The horizontal bars indicate the mean; e. Correlation between age and the percentage of images containing SNCs, SNCs with granules, and SNCs with granules and nucleolus. Groups are distinguished by color: black (control young), green (control), red (Parkinson), blue (Alzheimer). Each statistical test is a Mann-Whitney test with ns: not significant

### Identification of granule composition

While the histologists first considered lipofuscin accumulation in their evaluation, the much lower number of AF granules in age-matched control samples strongly argues against a purely age-related phenotype.

We therefore turned to staining with AmyT, an optical tracer that turns fluorescent when binding to β-sheet–rich structures. Targets include, but are not limited to amyloids or protein aggregates containing beta-amyloid fibrils and protofibrils. AmyT staining was present in pathological tissues and absent in controls ***(Fig.7a)***, comforting our hypothesis that AF inclusions were disease-specific amyloid-like deposits rather than lipofuscin inclusions ***(Fig.7b)***.

**Fig. 7.**
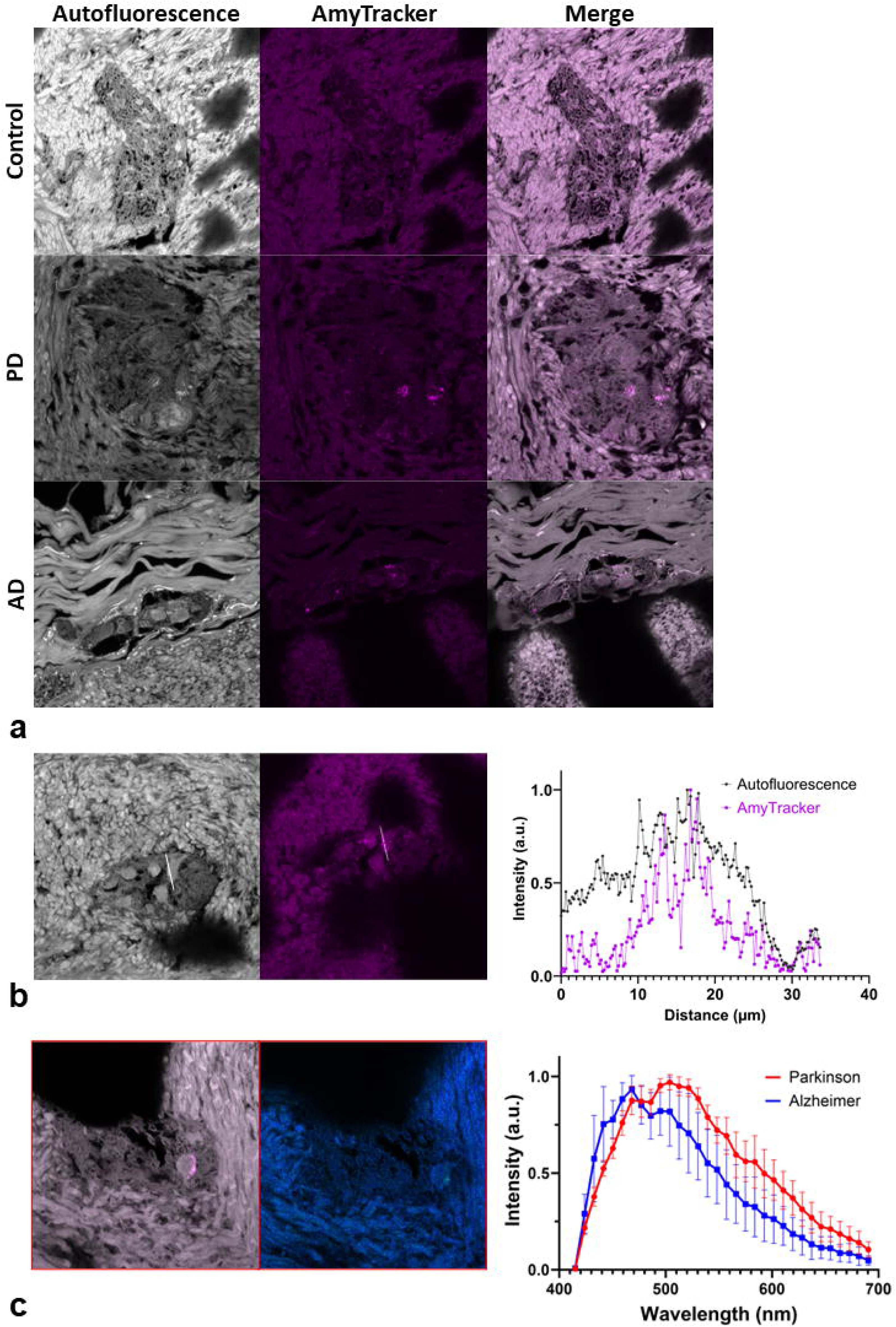
Identification of autofluorescent granules and analysis of their spectral properties. a. Representative images of AmyT labeling in control and pathological samples. AF is shown in grayscale. AmyT staining is shown in magenta; b. Colocalization analysis between AmyT labeling and AF granules; c. Spectral analysis of AF granules in PD (red) and AD (blue) samples

Spectral analysis revealed no significant differences between PD and AD granules ***(Fig.7c)***, indicating that AF granules distinguish pathological from control tissue, but they do not allow discriminating between disease types.

### Relationship Between LNC Presence and Diagnostic Accuracy in ENS Samples

By correlating our semi-quantitative analysis with the earlier blind diagnostic outcomes of the expert evaluation across almost 800 images, we confirmed a clear relationship between LNC presence and tissue (mis-) classification. Correctly diagnosed control samples were characterized by an absence or very low presence of LNCs, respectively, whereas controls misclassified as pathological frequently displayed LNCs, particularly those with cytoplasmic granules ***(Fig.8a)***. Conversely, correctly identified pathological samples showed a high prevalence of LNCs, especially granule-containing forms with or without a visible nucleolus ***(Fig.8b-c)***, while neuropathological samples misclassified as healthy typically exhibited only a few or no LNCs ***(Fig.8b-c)***. These results mirror our semi-quantitative findings, indicating that more than half of ENS ganglia in pathological samples may appear morphologically normal due to the absence of detectable LNCs.

**Fig. 8.**
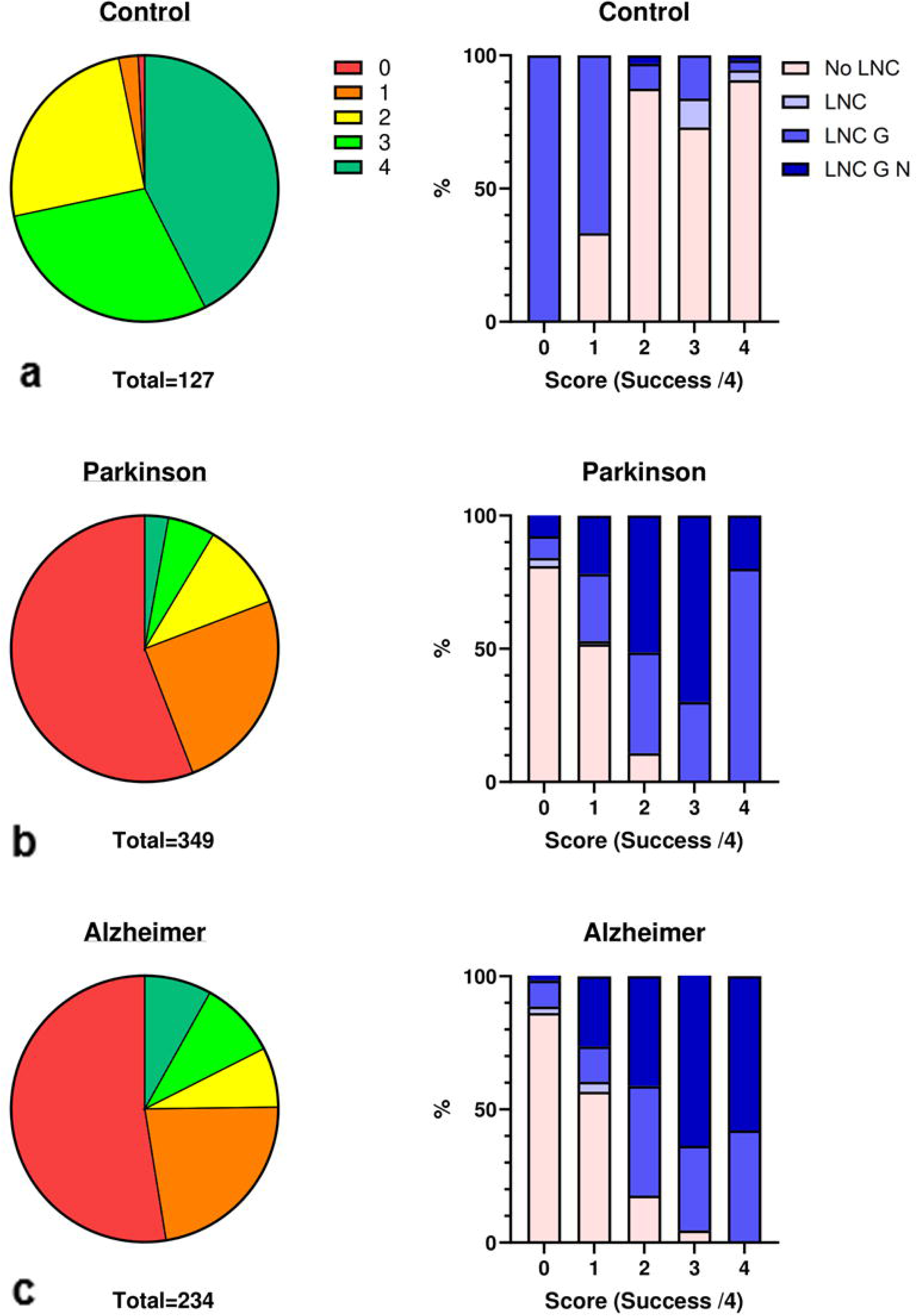
Relationship between the presence of LNC and blind diagnosis in ENS samples. a–c. Results of the blinded diagnosis based on AF images from healthy controls (a, n = 127), patients with Parkinson’s disease (b, n = 349), and patients with Alzheimer’s disease (c, n = 234); Left: pie charts illustrating the distribution of scores from 0 to 4, where 0 indicates that no expert histologist gave the correct diagnosis and 4 means that all expert histologists gave the correct diagnosis; Right: Stacked histograms showing, for each condition, the proportion of different types of LNCs for each score, with pink indicating no LNCs (No LNC) in the images, light blue indicating the presence of LNCs (LNC), blue indicating the presence of LNCs with granules (LNC G), and dark blue indicating the presence of LNCs with granules and nucleoli (LNC GN)

This heterogeneity reflects a fundamental limitation of ENS architecture, not methodological inconsistency. Because the ENS is a discontinuous, ganglionated network, diagnostically informative features are sparsely distributed and unlikely to be captured in limited 2D histological sections. As a result, confident diagnosis requires examining many ganglia per patient, which is often impractical in routine workflows.

### Sampling Limitations of 2D Sections and the Rationale for 3-D ENS Imaging

We therefore applied AF imaging to intact human colonoscopy biopsies. Owing to their small size and low pigmentation, no depigmentation or tissue clearing was required, preserving the fragile mucosal and submucosal architecture. PD biopsies were imaged uncleared to maintain tissue integrity ***(Fig.9a)***.

**Fig. 9.**
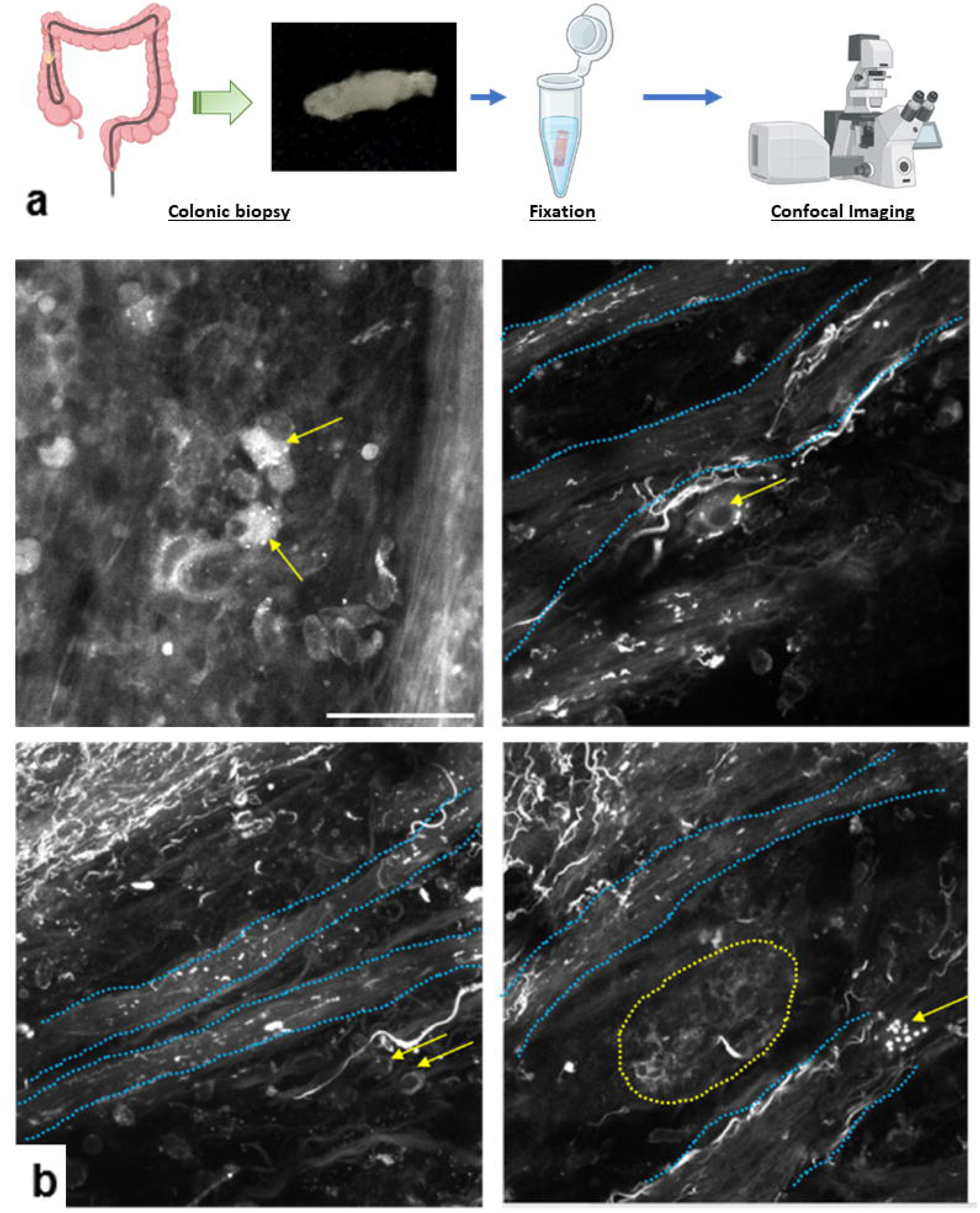
Autofluorence imaging of pathologic fixed human colon biopsies. a. Adaptation of the 3-D AF imaging workflow for thick samples. b. Representative images of nerve structures in the submucosal plexus obtained from colon biopsies. Yellow arrows: individual neurons of the submucosal plexus; blue dotted-lines: interganglionic nerve connections; yellow dotted-cercle: submucosal ganglion; scale bar: 50 µm. *(created with BioRender)*

As in thin sections, AF imaging enabled the visualization of submucosal structures ***(Fig.9b)***, including that of individual neurons within submucosal ganglia (yellow arrows). These cells displayed the same characteristic LNC morphology as observed previously, with cytoplasmic AF and brightly fluorescent granules, recapitulating our earlier findings from thin slices. In addition to identifying LNCs, we visualized interganglionic nerve connections (blue dotted lines), revealing the broader network organization of the ENS. In some cases, entire submucosal ganglia contained only neurons with a healthy morphology (yellow dotted outline), further supporting the idea that ENS pathology in PD is heterogeneous, with only a subset of ganglia affected.

Overall, our results demonstrate the feasibility of 3-D AF imaging on fixed, whole human colonic biopsies ***(Fig.SI3 – SI6)***. By providing a more complete and spatially integrated view of the ENS, biopsy-based volumetric imaging increases the likelihood of sampling a sufficient number of ganglia for reliable diagnosis. The observation of enteric neurons in PD biopsies exhibiting LNC-like morphology further supports the diagnostic relevance of our approach.

## Discussion

We established a large-scale label-free AF imaging workflow enabling the detailed analysis of ENS ganglia and cellular morphology from histological sections. The combination of slide-scanner wide-field acquisition with high-resolution confocal imaging of selected ROIs, facilitated by a coordinate-conversion macro, overcomes the limited field of view of confocal microscopy while preserving its spatial resolution.

Using this workflow, we generated a database of ∼800 confocal images of human myenteric and submucosal plexuses from controls and patients diagnosed with either PD or AD. While this dataset serves as a solid basis for morphometry and diagnostic applications it remains insufficient for robust deep-learning training, which will require several thousand annotated images.

AF imaging revealed an atypical neuronal phenotype characterized by hypertrophic cell bodies and variably containing cytoplasmic granules and/or a visible nucleolus. These LNCs were both observed in myenteric and submucosal plexuses from PD and AD cases but they were rare in controls, including aged individuals, suggesting that they are associated with neurodegenerative pathology rather than physiological aging. Their frequent presence in PD and AD supports their potential value as a marker of ENS neurodegeneration. Our observations are consistent with previous reports of enteric neuronal alterations in PD [47, 49], including the study by Gelpi et al. [26] from which these tissue sections were taken.

Together, our findings indicate that AF imaging is a robust approach for high-resolution morphological analysis of the ENS in neurodegenerative disorders, including PD and AD.

### Nature of AF granules in LNCs

The nature of the AF granules observed in LNCs remains an important question. Based on their spectral properties, lipofuscin was initially considered. However, lipofuscin accumulation is strongly age-dependent, so that similar deposits would be expected in age-matched controls. Their marked enrichment in pathological tissues argues against this interpretation. In contrast, AmyT labeling was associated with these structures in pathological samples, and it colocalized with AF granules, indicating the presence of β-sheet–rich material compatible with amyloid aggregates. Together, these findings support our interpretation that the AF signal in LNCs reflects pathological protein aggregation rather than age-related pigment accumulation, consistent with aggregated α-synuclein in PD and misfolded protein species in AD [2, 18].

Despite the presence of amyloid-like aggregates in both PD and AD, no disease-specific AF spectral or morphological signatures could be identified. AF imaging therefore distinguishes control from neuropathological ENS tissue through detection of protein aggregates, but it does not discriminate between neurodegenerative diseases. While this might reflect shared mechanisms of protein aggregation and impaired proteostasis across neurodegenerative disorders, it might also be due to the limited spectral resolution offered by the 32-element detector of our confocal microscope.

An important advantage of our approach is its applicability to older tissue samples. The present analyses were performed on the same sections used in the original study by Gelpi [26], allowing direct comparison with previously reported findings. However, the age of these samples limited the use of conventional immunofluorescence. This may reflect loss of antigenicity over time, incomplete information regarding previous tissue processing or storage conditions. Under these conditions, the combination of AF imaging and AmyT labeling provided the most robust approach to characterize these deposits. Beyond supporting the presence of aggregated protein material, this strategy highlights the value of AF-based imaging for the analysis of older ENS samples. It also suggests that tissue collections, even when no longer suitable for standard immunostaining, may still provide useful information on neuropathological changes.

### Putative identity and pathological significance of LNCs

Due to the impossibility of performing immunofluorescence on the old tissue sections, the cellular identity of LNCs remains unresolved. A dopaminergic phenotype appears unlikely given the low abundance of dopaminergic neurons in the human ENS (<5%) [34]. A cholinergic origin is more plausible, particularly within the myenteric plexus, where cholinergic neurons predominate and regulate intestinal motility under vagal control. Cholinergic pathways, including the ENS-vagal axis, are known to be selectively vulnerable in early PD [40, 46], supporting the hypothesis that LNCs might represent degenerating cholinergic neurons within a gut-brain axis process consistent with Braak’s model [6–10].

A subset of LNCs displaying both cytoplasmic granules and a prominent nucleolus was of particular interest. Increasing evidence implicates nucleolar dysfunction in the pathogenesis of both PD and AD through disruption of ribosomal biogenesis and activation of cellular stress pathways, including p53 signaling. In PD models, nucleolar impairment in dopaminergic neurons induces progressive neuronal loss and motor deficits [37, 41], and α-synuclein overexpression has also been linked to nucleolar stress [25]. Similarly, nucleolar alterations have been described in vulnerable neuronal populations in AD, notably in the hippocampus and entorhinal cortex, in association with tau and Aβ pathology [29, 32].

The presence of cytoplasmic amyloid-like aggregates together with nucleolar alterations in LNCs, and their absence in controls, suggests that these cells exhibit features of protein overload and nucleolar stress, two central mechanisms of neurodegeneration. These morphological characteristics may therefore represent cellular markers of neurodegenerative change in the ENS and could constitute early tissue biomarkers of enteric neuropathology.

### Diagnostic potential and limitations of ENS AF imaging

Blind diagnostic analyses highlighted a major limitation related to tissue heterogeneity. Pathological features are not uniformly present across ganglia, even within heavily affected cases, indicating that reliable assessment requires analysis of a sufficiently large number of ganglia. This observation supports the development of 3-D ENS imaging approaches on biopsies, which enables the simultaneous evaluation of many ganglia within a single volume acquisition.

Given the growing prevalence of neurodegenerative diseases and the absence of curative therapies, early diagnosis remains a major challenge [28, 38, 43]. Current biomarker strategies, including molecular assays, genetic approaches, brain imaging, and CSF analyses, are often costly, technically demanding, or difficult to implement at large scale [36, 50, 52]. In this context, ENS-based morphological label-free analysis of routine colonic biopsies offers a complementary and potentially accessible approach. Direct visualization of enteric neuronal alterations may provide disease-relevant structural information without relying on invasive or high-cost procedures.

Overall, the here presented simple and robust AF imaging workflow, compatible with existing workflows and standard clinical samples, provides a framework for the development of ENS-based biomarkers and may contribute to future early diagnostic strategies for neurodegenerative disorders.

**Fig.SI1 Questionnaire used for the blinded survey of histologists**

**Fig.SI2 Questionnaire used for the semi-quantitative study**

**Fig.SI3 – SI5 Animated autofluorescence z-stack of pathologic fixed human colon biopsies** Optical sections acquired every 2 µm; scale bar: 25 µm

**Fig.SI6 Animated autofluorescence z-stack of pathologic fixed human colon biopsies** Optical sections acquired every 5 µm; scale bar: 25 µm

## Supporting information

Fig.SI1 Questionnaire used for the blinded survey of histologists

Fig.SI2 Questionnaire used for the semi-quantitative study

Fig.SI3 Animated autofluorescence z-stack of pathologic fixed human colon biopsies

Fig.SI4 Animated autofluorescence z-stack of pathologic fixed human colon biopsies

Fig.SI5 Animated autofluorescence z-stack of pathologic fixed human colon biopsies

Fig.SI6 Animated autofluorescence z-stack of pathologic fixed human colon biopsies

## Acknowledgements

The imaging platform of the UMS BioMedTech facilities (Université Paris Cité, CNRS UMS2009, INSERM US36) is acknowledged for support. We are also grateful to Nolwen Rey for her precious help and valuable assistance.

## Author contributions

Doriane Hazart: Data curation, Investigation, Visualization, Writing - original draft. Marwa Moulzir: Writing - review & editing. Brigitte Delhomme: Review & editing. Pascal Derkinderen; Malvyne Rolli-Derkinderen; François Cossais; Peter H. Neckel: Investigation et Writing - review & editing. Martin Oheim; Clément Ricard: Funding acquisition, Project administration, Supervision, Validation, Visualization, Writing - review & editing.

## Data availability

The data that support the findings of this study are available from the corresponding author upon reasonable request

## Bibliography

1. Abbott RD, Petrovitch H, White LR, Masaki KH, Tanner CM, Curb JD, Grandinetti A, Blanchette PL, Popper JS, Ross GW (2001) Frequency of bowel movements and the future risk of Parkinson’s disease. Neurology 57:456–462. doi: 10.1212/wnl.57.3.456

2. Barrenschee M, Zorenkov D, Böttner M, Lange C, Cossais F, Scharf AB, Deuschl G, Schneider SA, Ellrichmann M, Fritscher-Ravens A, Wedel T (2017) Distinct pattern of enteric phospho-alpha-synuclein aggregates and gene expression profiles in patients with Parkinson’s disease. Acta Neuropathol Commun 5:1. doi: 10.1186/s40478-016-0408-2

3. Berg D, Postuma RB, Adler CH, Bloem BR, Chan P, Dubois B, Gasser T, Goetz CG, Halliday G, Joseph L, Lang AE, Liepelt-Scarfone I, Litvan I, Marek K, Obeso J, Oertel W, Olanow CW, Poewe W, Stern M, Deuschl G (2015) MDS research criteria for prodromal Parkinson’s disease. Mov Disord 30:1600–1611. doi: 10.1002/mds.26431

4. Berg D, Postuma RB, Bloem B, Chan P, Dubois B, Gasser T, Goetz CG, Halliday GM, Hardy J, Lang AE, Litvan I, Marek K, Obeso J, Oertel W, Olanow CW, Poewe W, Stern M, Deuschl G (2014) Time to redefine PD? Introductory statement of the MDS Task Force on the definition of Parkinson’s disease. Mov Disord 29:454–462. doi: 10.1002/mds.25844

5. Bloem BR, Okun MS, Klein C (2021) Parkinson’s disease. The Lancet 397:2284–2303. doi: 10.1016/S0140-6736(21)00218-X

6. Braak H, Del Tredici K (2008) Invited Article: Nervous system pathology in sporadic Parkinson disease. Neurology 70:1916–1925. doi: 10.1212/01.wnl.0000312279.49272.9f

7. Braak H, Del Tredici K (2017) Neuropathological Staging of Brain Pathology in Sporadic Parkinson’s disease: Separating the Wheat from the Chaff. Journal of Parkinson’s Disease 7:S71–S85. doi: 10.3233/JPD-179001

8. Braak H, Sastre M, Bohl JRE, de Vos RAI, Del Tredici K (2007) Parkinson’s disease: lesions in dorsal horn layer I, involvement of parasympathetic and sympathetic pre- and postganglionic neurons. Acta Neuropathol 113:421–429. doi: 10.1007/s00401-007-0193-x

9. Braak H, Tredici KD, Rüb U, de Vos RAI, Jansen Steur ENH, Braak E (2003) Staging of brain pathology related to sporadic Parkinson’s disease. Neurobiology of Aging 24:197–211. doi: 10.1016/S0197-4580(02)00065-9

10. Braak H, de Vos RAI, Bohl J, Del Tredici K (2006) Gastric α-synuclein immunoreactive inclusions in Meissner’s and Auerbach’s plexuses in cases staged for Parkinson’s disease-related brain pathology. Neuroscience Letters 396:67–72. doi: 10.1016/j.neulet.2005.11.012

11. Cersosimo MG, Benarroch EE (2012) Pathological correlates of gastrointestinal dysfunction in Parkinson’s disease. Neurobiol Dis 46:559–564. doi: 10.1016/j.nbd.2011.10.014

12. Chen R, Gu X, Wang X (2022) α-Synuclein in Parkinson’s disease and advances in detection. Clin Chim Acta 529:76–86. doi: 10.1016/j.cca.2022.02.006

13. Corbillé A-G, Letournel F, Kordower JH, Lee J, Shanes E, Neunlist M, Munoz DG, Derkinderen P, Beach TG (2016) Evaluation of alpha-synuclein immunohistochemical methods for the detection of Lewy-type synucleinopathy in gastrointestinal biopsies. Acta Neuropathologica Communications 4:35. doi: 10.1186/s40478-016-0305-8

14. Costa M, Brookes SJH, Hennig GW (2000) Anatomy and physiology of the enteric nervous system. Gut 47:iv15–iv19. doi: 10.1136/gut.47.suppl_4.iv15

15. Dauer W, Przedborski S (2003) Parkinson’s Disease: Mechanisms and Models. Neuron 39:889–909. doi: 10.1016/S0896-6273(03)00568-3

16. Dickson DW (2018) Neuropathology of Parkinson disease. Parkinsonism & Related Disorders 46:S30–S33. doi: 10.1016/j.parkreldis.2017.07.033

17. Donadio V, Incensi A, Leta V, Giannoccaro MP, Scaglione C, Martinelli P, Capellari S, Avoni P, Baruzzi A, Liguori R (2014) Skin nerve α-synuclein deposits. Neurology 82:1362–1369. doi: 10.1212/WNL.0000000000000316

18. Dougnon G, Matsui H (2025) Lipofuscin accumulation in aging and neurodegeneration: a potential “timebomb” overlooked in Alzheimer’s disease. Transl Neurodegener 14:67. doi: 10.1186/s40035-025-00529-x

19. Du T, Wang L, Liu W, Zhu G, Chen Y, Zhang J (2021) Biomarkers and the Role of α-Synuclein in Parkinson’s Disease. Front Aging Neurosci 13:645996. doi: 10.3389/fnagi.2021.645996

20. Fairfoul G, McGuire LI, Pal S, Ironside JW, Neumann J, Christie S, Joachim C, Esiri M, Evetts SG, Rolinski M, Baig F, Ruffmann C, Wade-Martins R, Hu MTM, Parkkinen L, Green AJE (2016) Alpha-synuclein RT-QuIC in the CSF of patients with alpha-synucleinopathies. Annals of Clinical and Translational Neurology 3:812–818. doi: 10.1002/acn3.338

21. Fasano A, Visanji NP, Liu LWC, Lang AE, Pfeiffer RF (2015) Gastrointestinal dysfunction in Parkinson’s disease. The Lancet Neurology 14:625–639. doi: 10.1016/S1474-4422(15)00007-1

22. Fearnley JM, Lees AJ (1991) Ageing and Parkinson’s disease: substantia nigra regional selectivity. Brain 114 (Pt 5):2283–2301. doi: 10.1093/brain/114.5.2283

23. Furness JB (2012) The enteric nervous system and neurogastroenterology. Nat Rev Gastroenterol Hepatol 9:286–294. doi: 10.1038/nrgastro.2012.32

24. Ganguly U, Singh S, Pal S, Prasad S, Agrawal BK, Saini RV, Chakrabarti S (2021) Alpha-Synuclein as a Biomarker of Parkinson’s Disease: Good, but Not Good Enough. Front Aging Neurosci 13:702639. doi: 10.3389/fnagi.2021.702639

25. Garcia-Esparcia P, Hernández-Ortega K, Koneti A, Gil L, Delgado-Morales R, Castaño E, Carmona M, Ferrer I (2015) Altered machinery of protein synthesis is region- and stage-dependent and is associated with α-synuclein oligomers in Parkinson’s disease. Acta Neuropathologica Communications 3:76. doi: 10.1186/s40478-015-0257-4

26. Gelpi E, Navarro-Otano J, Tolosa E, Gaig C, Compta Y, Rey MJ, Martí MJ, Hernández I, Valldeoriola F, Reñé R, Ribalta T (2014) Multiple organ involvement by alpha-synuclein pathology in Lewy body disorders. Movement Disorders 29:1010–1018. doi: 10.1002/mds.25776

27. de Guilhem de Lataillade A, Caillaud M, Oullier T, Naveilhan P, Pellegrini C, Tolosa E, Neunlist M, Rolli-Derkinderen M, Gelpi E, Derkinderen P (2023) LRRK2 expression in normal and pathologic human gut and in rodent enteric neural cell lines. J Neurochem 164:193–209. doi: 10.1111/jnc.15704

28. Hampel H, O’Bryant SE, Molinuevo JL, Zetterberg H, Masters CL, Lista S, Kiddle SJ, Batrla R, Blennow K (2018) Blood-based biomarkers for Alzheimer disease: mapping the road to the clinic. Nat Rev Neurol 14:639–652. doi: 10.1038/s41582-018-0079-7

29. Hernández-Ortega K, Garcia-Esparcia P, Gil L, Lucas JJ, Ferrer I (2015) Altered Machinery of Protein Synthesis in Alzheimer’s: From the Nucleolus to the Ribosome. Brain Pathol 26:593–605. doi: 10.1111/bpa.12335

30. Houser MC, Tansey MG (2017) The gut-brain axis: is intestinal inflammation a silent driver of Parkinson’s disease pathogenesis? npj Parkinson’s Disease 3:3. doi: 10.1038/s41531-016-0002-0

31. Hughes AJ, Daniel SE, Ben-Shlomo Y, Lees AJ (2002) The accuracy of diagnosis of parkinsonian syndromes in a specialist movement disorder service. Brain 125:861–870. doi: 10.1093/brain/awf080

32. Koren SA, Galvis-Escobar S, Abisambra JF (2020) Tau-mediated dysregulation of RNA: Evidence for a common molecular mechanism of toxicity in frontotemporal dementia and other tauopathies. Neurobiology of Disease 141:104939. doi: 10.1016/j.nbd.2020.104939

33. Lebouvier T, Neunlist M, Varannes SB des, Coron E, Drouard A, N’Guyen J-M, Chaumette T, Tasselli M, Paillusson S, Flamand M, Galmiche J-P, Damier P, Derkinderen P (2010) Colonic Biopsies to Assess the Neuropathology of Parkinson’s Disease and Its Relationship with Symptoms. PLOS ONE 5:e12728. doi: 10.1371/journal.pone.0012728

34. Li ZS, Pham TD, Tamir H, Chen JJ, Gershon MD (2004) Enteric Dopaminergic Neurons: Definition, Developmental Lineage, and Effects of Extrinsic Denervation. J Neurosci 24:1330–1339. doi: 10.1523/JNEUROSCI.3982-03.2004

35. Magalhães P, Lashuel HA (2022) Opportunities and challenges of alpha-synuclein as a potential biomarker for Parkinson’s disease and other synucleinopathies. NPJ Parkinsons Dis 8:93. doi: 10.1038/s41531-022-00357-0

36. Olsson B, Lautner R, Andreasson U, Öhrfelt A, Portelius E, Bjerke M, Hölttä M, Rosén C, Olsson C, Strobel G, Wu E, Dakin K, Petzold M, Blennow K, Zetterberg H (2016) CSF and blood biomarkers for the diagnosis of Alzheimer’s disease: a systematic review and meta-analysis. The Lancet Neurology 15:673–684. doi: 10.1016/S1474-4422(16)00070-3

37. Parlato R, Liss B (2018) Selective degeneration of dopamine neurons in Parkinson’s disease: emerging roles of altered calcium homeostasis and nucleolar function. e-Neuroforum 24:A1–A9. doi: 10.1515/nf-2017-A006

38. Poewe W, Seppi K, Tanner CM, Halliday GM, Brundin P, Volkmann J, Schrag A-E, Lang AE (2017) Parkinson disease. Nat Rev Dis Primers 3:1–21. doi: 10.1038/nrdp.2017.13

39. Postuma RB, Berg D, Stern M, Poewe W, Olanow CW, Oertel W, Obeso J, Marek K, Litvan I, Lang AE, Halliday G, Goetz CG, Gasser T, Dubois B, Chan P, Bloem BR, Adler CH, Deuschl G (2015) MDS clinical diagnostic criteria for Parkinson’s disease. Movement Disorders 30:1591–1601. doi: 10.1002/mds.26424

40. Rao M, Gershon MD (2016) The bowel and beyond: the enteric nervous system in neurological disorders. Nat Rev Gastroenterol Hepatol 13:517–528. doi: 10.1038/nrgastro.2016.107

41. Rieker C, Engblom D, Kreiner G, Domanskyi A, Schober A, Stotz S, Neumann M, Yuan X, Grummt I, Schütz G, Parlato R (2011) Nucleolar Disruption in Dopaminergic Neurons Leads to Oxidative Damage and Parkinsonism through Repression of Mammalian Target of Rapamycin Signaling. J Neurosci 31:453–460. doi: 10.1523/JNEUROSCI.0590-10.2011

42. Russo MJ, Orru CD, Concha-Marambio L, Giaisi S, Groveman BR, Farris CM, Holguin B, Hughson AG, LaFontant D-E, Caspell-Garcia C, Coffey CS, Mollon J, Hutten SJ, Merchant K, Heym RG, Soto C, Caughey B, Kang UJ (2021) High diagnostic performance of independent alpha-synuclein seed amplification assays for detection of early Parkinson’s disease. Acta Neuropathologica Communications 9:179. doi: 10.1186/s40478-021-01282-8

43. Scheltens P, Strooper BD, Kivipelto M, Holstege H, Chételat G, Teunissen CE, Cummings J, Flier WM van der (2021) Alzheimer’s disease. The Lancet 397:1577–1590. doi: 10.1016/S0140-6736(20)32205-4

44. Schindelin J, Arganda-Carreras I, Frise E, Kaynig V, Longair M, Pietzsch T, Preibisch S, Rueden C, Saalfeld S, Schmid B, Tinevez J-Y, White DJ, Hartenstein V, Eliceiri K, Tomancak P, Cardona A (2012) Fiji: an open-source platform for biological-image analysis. Nat Methods 9:676–682. doi: 10.1038/nmeth.2019

45. Shannon KM, Keshavarzian A, Mutlu E, Dodiya HB, Daian D, Jaglin JA, Kordower JH (2012) Alpha-synuclein in colonic submucosa in early untreated Parkinson’s disease. Movement Disorders 27:709–715. doi: 10.1002/mds.23838

46. Shprecher DR, Derkinderen P (2012) Parkinson disease: the enteric nervous system spills its guts. Neurology 78:683; author reply 683. doi: 10.1212/WNL.0b013e31824bd195

47. Singaram C, Ashraf W, Gaumnitz EA, Torbey C, Sengupta A, Pfeiffer R, Quigley EM (1995) Dopaminergic defect of enteric nervous system in Parkinson’s disease patients with chronic constipation. Lancet 346:861–864. doi: 10.1016/s0140-6736(95)92707-7

48. Stirpe P, Hoffman M, Badiali D, Colosimo C (2016) Constipation: an emerging risk factor for Parkinson’s disease? European Journal of Neurology 23:1606–1613. doi: 10.1111/ene.13082

49. Stokholm MG, Danielsen EH, Hamilton-Dutoit SJ, Borghammer P (2016) Pathological α-synuclein in gastrointestinal tissues from prodromal Parkinson disease patients. Annals of Neurology 79:940–949. doi: 10.1002/ana.24648

50. Villemagne VL, Doré V, Burnham SC, Masters CL, Rowe CC (2018) Imaging tau and amyloid-β proteinopathies in Alzheimer disease and other conditions. Nat Rev Neurol 14:225–236. doi: 10.1038/nrneurol.2018.9

51. Wang Z, Becker K, Donadio V, Siedlak S, Yuan J, Rezaee M, Incensi A, Kuzkina A, Orrú CD, Tatsuoka C, Liguori R, Gunzler SA, Caughey B, Jimenez-Capdeville ME, Zhu X, Doppler K, Cui L, Chen SG, Ma J, Zou W-Q (2021) Skin α-Synuclein Aggregation Seeding Activity as a Novel Biomarker for Parkinson Disease. JAMA Neurology 78:30–40. doi: 10.1001/jamaneurol.2020.3311

52. Zetterberg H, Blennow K (2021) Moving fluid biomarkers for Alzheimer’s disease from research tools to routine clinical diagnostics. Molecular Neurodegeneration 16:10. doi: 10.1186/s13024-021-00430-x

53. Overview of the Digestive System | Anatomy and Physiology II. https://courses.lumenlearning.com/suny-ap2/chapter/overview-of-the-digestive-system/?utm_source=chatgpt.com. Accessed 1 July 2025

