## Supplementary figures and images for "Diagnosing Neurodegenerative Diseases by Label-Free 3-D Imaging of Intestinal Samples"

### Fig.SI1 Questionnaire used for the blinded survey of histologists

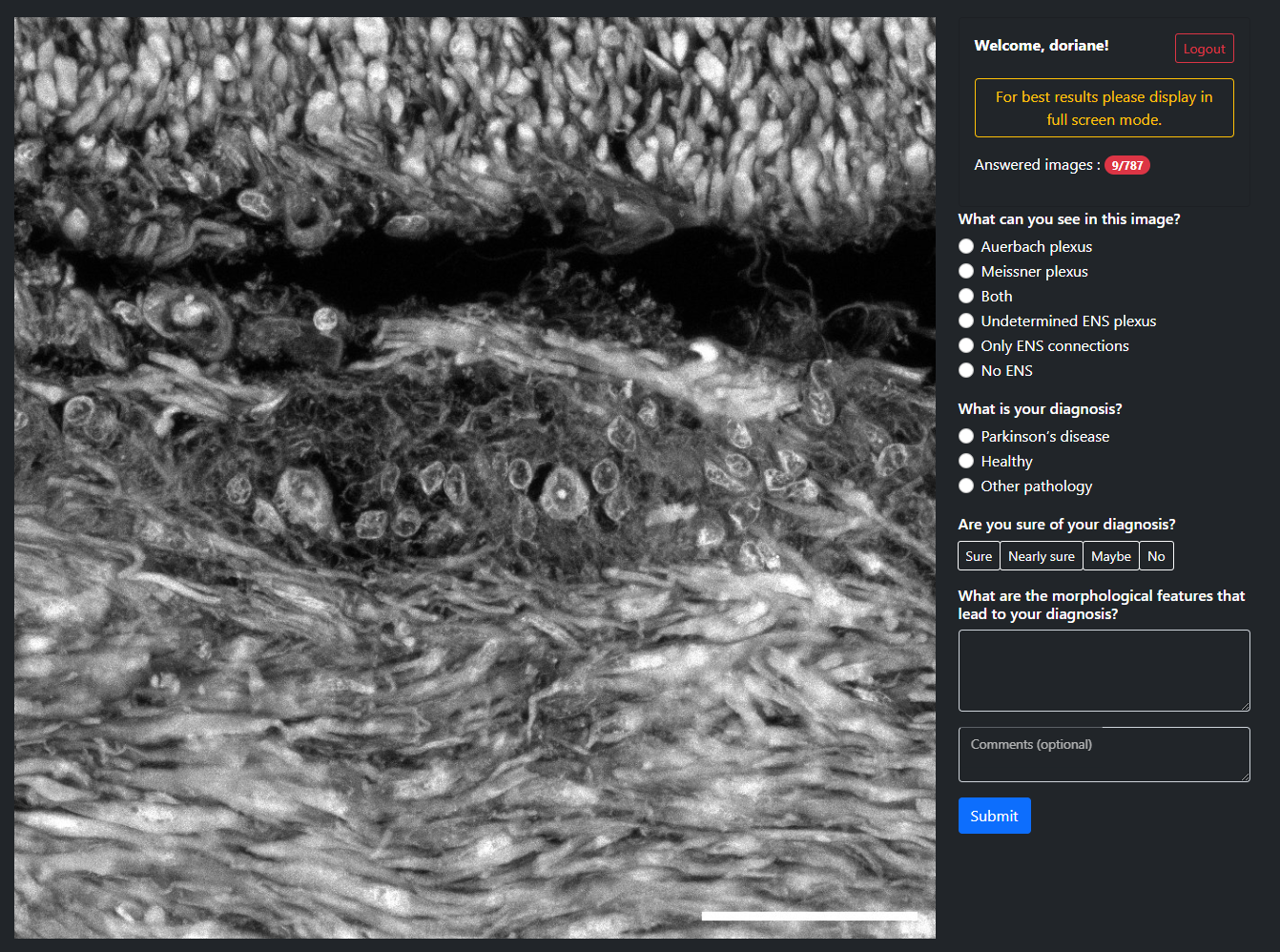

### Fig.SI2 Questionnaire used for the semi-quantitative study

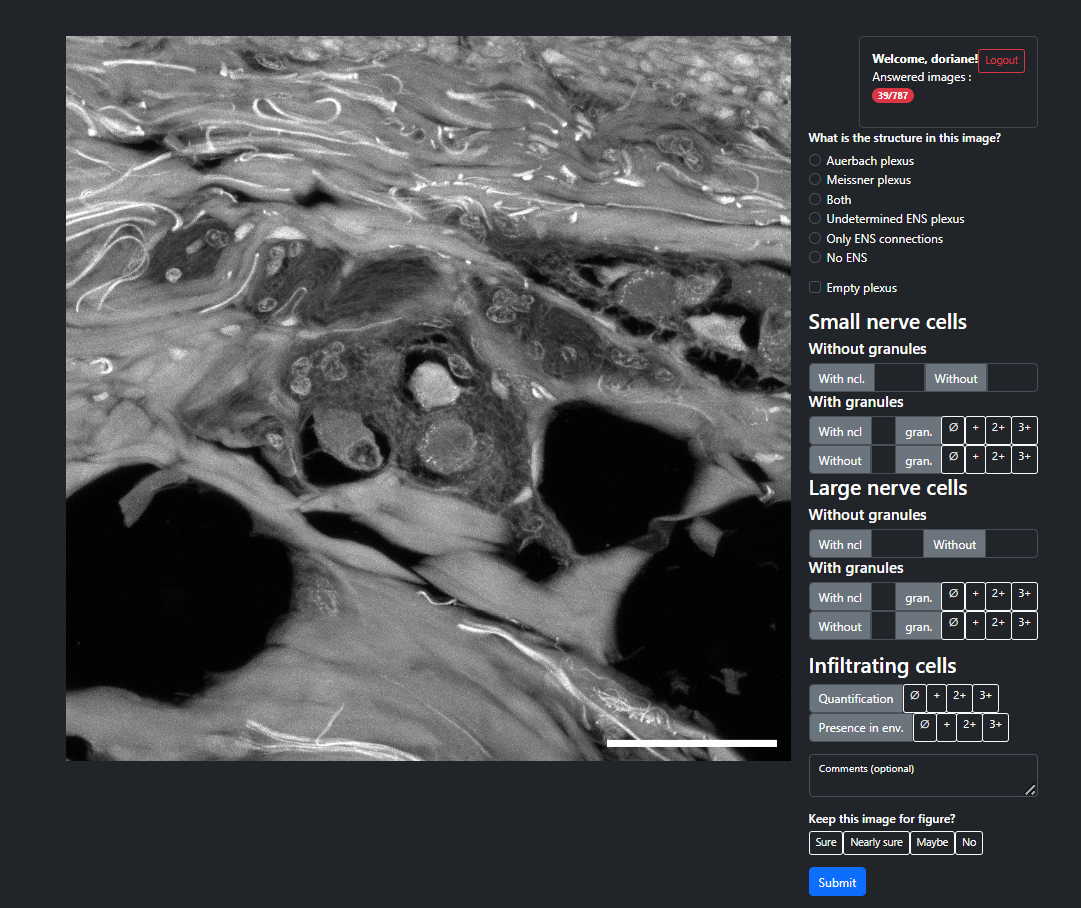
